# Global post-translational modification profiling of HIV-1-infected cells reveals mechanisms of host cellular pathway remodeling

**DOI:** 10.1101/2020.01.06.896365

**Authors:** Jeffrey R. Johnson, David C. Crosby, Judd F. Hultquist, Donna Li, John Marlett, Justine Swann, Ruth Hüttenhain, Erik Verschueren, Tasha L. Johnson, Billy W. Newton, Michael Shales, Pedro Beltrao, Alan D. Frankel, Alexander Marson, Oliver I. Fregoso, John A. T. Young, Nevan J. Krogan

## Abstract

Viruses must effectively remodel host cellular pathways to replicate and evade immune defenses, and they must do so with limited genomic coding capacity. Targeting post-translational modification (PTM) pathways provides a mechanism by which viruses can broadly and rapidly transform a hostile host environment into a hospitable one. We used quantitative proteomics to measure changes in two PTM types – phosphorylation and ubiquitination – in response to HIV-1 infection with viruses harboring targeted deletions of a subset of HIV-1 genes. PTM analysis revealed a requirement for Aurora kinase A activity in HIV-1 infection and furthermore revealed that AMP-activated kinase activity is modulated during infection via HIV-1 Vif-mediated degradation of B56-containing protein phosphatase 2A (PP2A). Finally, we demonstrated that the Cullin4A-DDB1-DCAF1 E3 ubiquitin ligase ubiquitinates histone H1 somatic isoforms and that HIV-1 Vpr inhibits this process, leading to defects in DNA repair. Thus, global PTM profiling of infected cells serves as an effective tool for uncovering specific mechanisms of host pathway modulation.

## INTRODUCTION

Human immunodeficiency virus type-1 (HIV-1) remains a major threat to public health worldwide. While there has been tremendous success in the development of combination antiretroviral therapies (cART) for treatment and prophylaxis, there is a need for next-generation therapies that are less toxic, have fewer side effects, and are longer-acting. Drug resistance is also a threat to existing therapies, with an estimated 10 percent of people received cART that are resistant to at least one drug in their cART (Organization, 2017. The identification of new drug targets is critical to next-generation antiretroviral drug development, and as such a precise molecular understanding of HIV-1 gene functions is of critical importance.

The HIV-1 genome encodes several accessory genes that are dispensable for replication in some cell lines but are required for evasion of host antiviral pathways and in vivo replication and pathogenesis. Three accessory genes, *Vif*, *Vpr*, and *Vpu*, bind to Cullin RING-type E3 ubiquitin ligases in order to rewire host ubiquitination systems and antagonize antiviral host pathways (Sauter and Kirchhoff, 2018) binds to the Cullin-5/Elogin-B/Elongin-C RING-type E3 ubiquitin ligase (CRL5^EloB/EloC^) in order to promote ubiquitination and proteasomal degradation of antiviral APOBEC3 proteins, a family of single-stranded DNA-editing enzymes with the capacity to incorporate into HIV-1 virions and inactivate the virus by hypermutation of the genome during reverse transcription (eehy et al., 2002; Harris et al., 2003). Vif also targets the PP2A-B56 subfamily of protein phosphatases for ubiquitination and degradation, and this degradation is associated with G2 cell cycle arrest (Greenwood et al., 2016; Salamango et al., 2019). The substrate(s) of PP2A-B56 affected by its degradation and the molecular mechanism underlying Vif-mediated G2 arrest remains unknown. The HIV-1 Vpu protein binds to the Cullin-1/Skp1/βTrCP ligase (SCF^βTrCP^) to promote ubiquitination and removal from the cell surface of BST/tetherin, an antiviral protein that inhibits viral budding and release (Neil et al., 2008). HIV-2, an independent zoonotic transmission of simian immunodeficiency virus (SIV) from primate in humans that is distinct from HIV-1, encodes the additional Vpx protein that binds to the Cullin4/DDB1/DCAF1 (CRL4^DCAF1^) ligase to promote ubiquitination and degradation of the antiviral protein SAMHD1, which restricts free deoxynucleotide pools and impairs viral reverse transcription (Hrecka et al., 2011; Laguette et al., 2011) Intriguingly, HIV-2 Vpx also utilizes CRL4^DCAF1^ to promote ubiquitination and degradation of the HUSH transcriptional silencing complex that otherwise acts to silence incoming proviruses in the nucleus (Chougui et al., 2018; Dahabieh et al., 2015; Lai and Pugh, 2017; Yurkovetskiy et al., 2018). It is not yet clear whether HUSH is wired to the innate immune system. The HIV-1 Vpr protein also binds to CRL4^DCAF1^, but it does not share SAMHD1 and HUSH inactivation functions with HIV-2 Vpx. Several Vpr-CRL4^DCAF1^ substrates have been identified including the uracil N-glycosylase 2 (UNG2), Mus81 (part of the SLX4 complex), helicase-like transcription factor (HLTF), exonuclease 1 (EXO1), and Tet DNA dioxygenase 2 (TET2) (Laguette et al., 2014; Lahouassa et al., 2016; Lv et al., 2018; Yan et al., 2018. However, only TET2 has been demonstrated to rescue a replication defect of Vpr-deficient viral replication in primary monocyte-derived macrophages and the rescue was only partial (Wang and Su, 2019) Furthermore, none of the Vpr-CRL4^DCAF1^ substrates identified thus far explain the ability of Vpr to potently activate cell cycle arrest at the G2/M-phase checkpoint.

The identification of post-translational modification (PTM) enzyme-substrate relationships can be challenging due to the relatively transient nature of their physical interactions. The discoveries of APOBEC3 and BST2/tetherin proteins as ubiquitination targets of Vif-CRL5^ELOB/ELOC^ and Vpu-SCF^βTrCP^, respectively, were made by comparing gene expression patterns in closely related cell lines that were permissive or non-permissive to HIV-1 replication with Vif- and Vpu-deficient viruses. Conversely, the suite of Vpr and Vpx substrates have been identified by physical interactions with Vpr/Vpx-CRL4^DCAF1^, as suitable non-permissive cell lines for HIV-1 replication with Vpr- or Vpx-deficient viruses have not been established. Given that none of the Vpr-CRL4^DCAF1^ substrates identified thus far explains its ability to arrest the cell cycle, it is likely that additional Vpr-CRL4^DCAF1^ substrate(s) remain to be discovered.

The development of chemical and immunoaffinity methods to enrich for ubiquitinated and phosphorylated species, combined with increasingly sensitive and comprehensive quantitative mass spectrometry-based proteomics approaches, allows for a “shotgun” approach to identify PTM enzyme-substrate relationships. In this study, we sought to identify PTM pathways perturbed by HIV-1 infection in order to better understand the molecular mechanisms underlying HIV-1 accessory gene functions. We applied a global mass spectrometry-based proteomics approach to obtain a quantitative survey of two major PTM types – ubiquitination and phosphorylation – in cells infected with wild-type and accessory gene-deficient HIV-1 viruses. These data represent a quantitative resource of PTM changes during HIV-1 infection that will facilitate future investigations of the relationship between HIV-1 and its host.

## RESULTS

### Proteome-wide evaluation of ubiquitination responses to HIV-1 infection

Since there are clear mechanisms of ubiquitination machinery hijacking by HIV-1 we first sought to comprehensively quantify changes in ubiquitination in response to HIV-1 infection. To do so we combined a stable isotope labeling of amino acids in culture (SILAC) approach with ubiquitin remnant purification and mass spectrometry (Figure 1A) (Ong et al., 2002). Briefly, SILAC-labeled Jurkat T cells were infected with *Env*-deficient HIV-1 (strain NL4-3, referred to as WT hereafter) that was pseudotyped with vesicular stomatitis virus glycoprotein (VSVG) at a multiplicity of infection of 5 to achieve infection of >95% of cells. Infected cells were compared to mock-infected cells treated with *Env*-deficient HIV-1 without VSVG-pseudotyping to prevent cell entry. Infections were performed in biological quadruplicate with SILAC labels inverted across experiments. In order to stabilize ubiquitinated substrates that may be rapidly degraded by the proteasome, all ubiquitination experiments were performed in both the presence and absence of a proteasome inhibitor, MG-132, provided at 10 μ harvesting. We have previously demonstrated that this treatment is sufficient to stabilize ubiquitinated peptides derived from canonical HIV-1 ubiquitination targets APOBEC3C and CD4 during infection (Ball et al., 2016). 24 hours post-infection, cells were lysed and lysates derived from HIV-1- and mock-infected cells were combined at equal protein concentrations and subjected to trypsin digestion, ubiquitin remnant immunoaffinity enrichment, and quantitative mass spectrometry analysis on an Orbitrap Elite mass spectrometry system (Xu et al., 2010). An aliquot of trypsin-digested material prior to ubiquitin remnant immunoaffinity enrichment was reserved for hydrophilic interaction chromatography (HILIC) fractionation and protein abundance analysis. Raw mass spectrometry data was first analyzed by MaxQuant to identify ubiquitinated peptides, localize ubiquitination sites, and extract SILAC ratios and ion intensities (Cox and Mann, 2008). Subsequently, the MSstats statistical package was employed to model variability at all levels (*i.e.*, biological variability, technical variability, and variability at the level of peptides, proteins, and treatment or infection conditions), and to apply a statistical tests to calculate the probability that ubiquitination changes could be explained by the model of variation, and to adjust for multiple testing (Choi et al., 2014).

**Figure 1.**
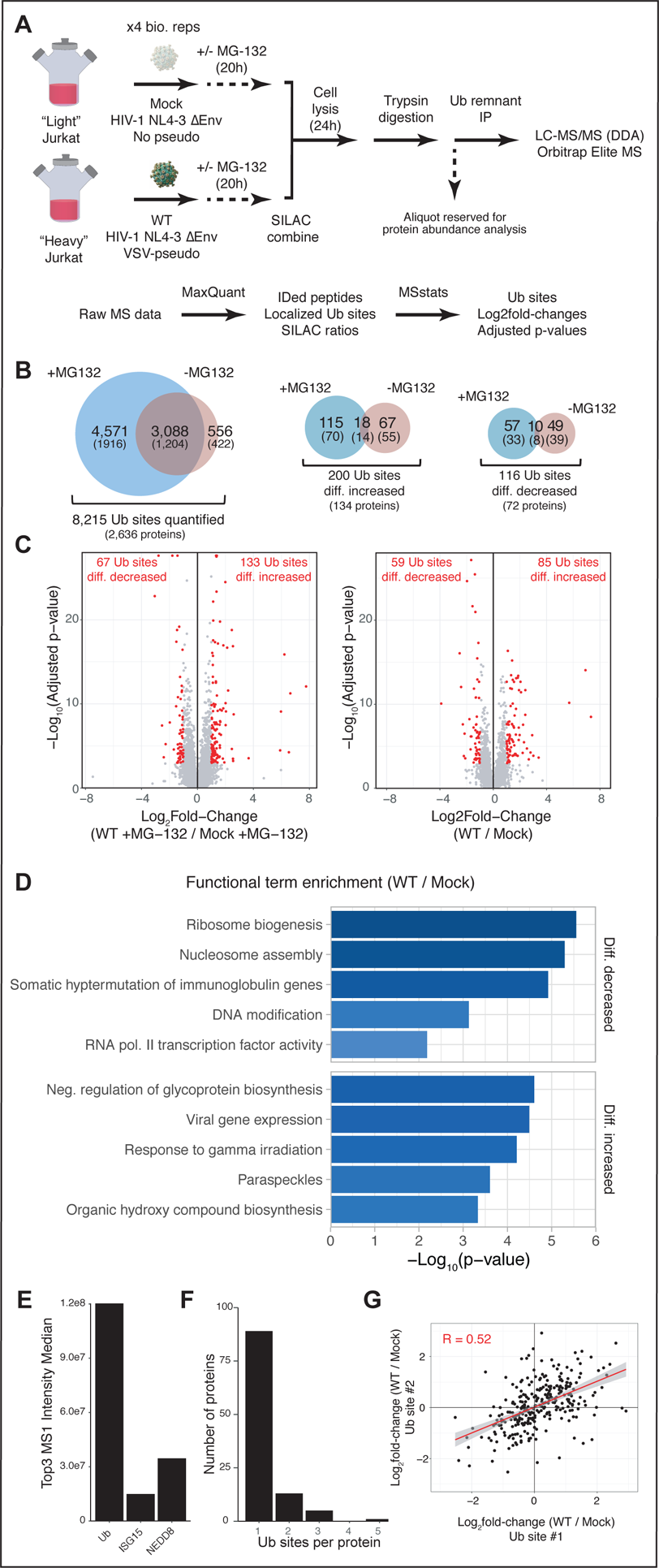
**Proteome-wide evaluation of ubiquitination responses to HIV-1 infection.** (A) Schematic of experimental and data analysis workflows implemented (B) Overlaps of ubiquitination sites detected and differentially increased or decreased by HIV-1 infection in the absence and presence of proteasome inhibitor (C) Volcano plots of ubiquitination changes in the presence and absence of proteasome inhibitor, MG-132. Ubiquitination sites considered differentially increased or decreased with |log_2_fold-change| > 1.0 and adjusted p-value < 0.05 are highlighted in red. (D) Pathway and process enrichment of HIV-1-regulated ubiquitination sites (E) Top3 MS1 intensity estimation of absolute protein abundance for ubiquitin and ubiquitin-like proteins that yield diglycine remnants upon trypsin digestion (F) Number of differentially increased or decreased ubiquitination sites per protein (G) Scatterplot of log_2_fold-changes for consecutive ubiquitination sites on the same protein

7,659 and 3,644 ubiquitination sites on 3,120 and 1,626 proteins were identified in experiments performed in the presence and absence of proteasome inhibitor, respectively, with 8,325 ubiquitination sites on 2,636 proteins identified in total (Figure 1B, Supplementary Table S1). 85% of ubiquitination sites (3,088) detected in the absence of proteasome inhibitor were also detected in its presence (Figure 1B).

Proteasome inhibition increased the number of sites identified by over 2-fold. We considered ubiquitination sites to be differentially decreased or increased if they were observed with log_2_fold-change (WT/mock) < −1 or log_2_fold-change (WT/mock) > 1, respectively, with an adjusted p-value < 0.05 (Figure 1C). By these criteria, the overlap of ubiquitination sites that were differentially increased or decreased in both the presence and absence of proteasome inhibitor was 21% (18 sites) and 17% (10 sites), respectively. While proteasome inhibition was required to observe HIV-1-mediated ubiquitination on CD4 and APOBEC3C – both of which are ubiquitinated and targeted for proteasomal degradation by HIV-1 – proteasome inhibition has also been demonstrated to antagonize HIV-1 replication and is likely to impact or even reverse many HIV-1-mediated ubiquitination changes (Miller et al., 2013). The low overlap between ubiquitination sites that were observed to be differentially abundant in the presence and absence of proteasome inhibition likely reflects the profound effects that proteasome inhibition has on ubiquitination profiles in HIV-1-infected cells.

We next performed pathway and process enrichment analysis for proteins with ubiquitination sites that were differentially decreased or increased during HIV-1 infection using the Metascape tool (Figure 1D) (Zhou et al., 2019) inhibition had profound effects on ubiquitination profiles in HIV-infected cells, we restricted this analysis to ubiquitination profiles from samples that were not proteasome inhibited. This enrichment analysis uncovered differentially decreased ubiquitination of proteins involved in nucleosome assembly (genes HIST1H1C, HIST1H1D, HIST1H1B,

H1FX, MIS18A, HJURP; p-value = 5.07 x 10^-6^), which included four histone H1 variants that were less ubiquitinated in infected cells. Enrichment analysis also uncovered differentially increased ubiquitination of proteins involved in the response to gamma irradiation (genes PARP1, ATR, CBL, FANCD2, XRCC6, H2AFX, XRCC5, p-value = 6.19×10^-5^), which may be a result of HIV-1-mediated activation of the DNA damage response pathway via the *Vpr* gene (Bartz et al., 1996; Roshal et al., 2003)

Ubiquitin is not the only protein known to generate a diglycine ubiquitin remnant. Ubiquitin-like proteins ISG15 and NEDD8, which are also covalently attached the substrate lysine residues, also generate a diglycine remnant upon trypsin digestion. We employed a ‘Top 3’ intensity average method to estimate the relative abundance of ubiquitin, ISG15, and NEDD8 from the protein abundance analysis of these lysates (Figure 1E) (Grossmann et al., 2010). By this method we estimate that ubiquitin is 5-10 times more abundant than ISG15 and NEDD8, and thus we expect most sites discovered in our study to be ubiquitin modifications.

Many proteins were observed with multiple ubiquitination sites that were similarly regulated by HIV-1 infection (Figure 1F). To analyze the behavior of ubiquitination sites on multiply ubiquitinated proteins we plotted the log_2_fold-change values for consecutive ubiquitination sites on the same protein for proteins with at least two regulated ubiquitination sites (Figure 1G). Overall, ubiquitination sites on the same multiply ubiquitinated proteins were correlated with a Pearson coefficient of 0.52, with 75% of ubiquitination sites changing in the same direction as other ubiquitination sites on the same protein. Of the proteins that were differentially ubiquitinated at multiple residues, cleavage and polyadenylation specificity factors 5 and 6 (CPSF5 and CPSF6), members of the cleavage factor I 3’ end processing complex, were observed with differentially increased ubiquitination at 3 and 2 sites, respectively. CPSF6 has been demonstrated to interact with the HIV-1 capsid and plays an important role in nuclear entry and HIV-1 integration site targeting (Bejarano et al., 2019; Lee et al., 2010; Sowd et al., 2016. Similarly, XRCC5/Ku80 and XRCC6/Ku70, which comprise the Ku complex that is 2016) required for non-homologous end-joining (NHEJ) DNA repair, were each observed with differentially increased ubiquitination at two different sites.

To identify ubiquitination sites that may impact protein activity or function, we compared the positions of differentially abundant ubiquitination sites with UniProt protein domain definitions to identify 47 regulated ubiquitination sites that occurred within protein domains (Figure 2A) (Bateman et al., 2018). This analysis also identified differentially increased ubiquitination sites on the Ku complex that occurred within the Ku domain of XRCC5/Ku80 and the SAP domain of XRCC6/Ku70 (Figure 2B) (Walker et al., 2001) We hypothesized that Ku may act as a restriction factor that limits HIV-1 infection and that this activity is counteracted by HIV-1-mediated ubiquitination. To test the capacity of Ku to act as an HIV-1 restriction factor we utilized a CRIPSR/Cas9 nucleofection approach to genetically perturb Ku in human primary CD4+ T cells and to test the effects (Hultquist et al., 2019. Using two crRNAs targeting the XRCC5/Ku80 locus we confirmed reduction of XRCC5/Ku80 protein levels by Western blot and observed an increase in HIV-1 infection in three independent donors that was up to 3-fold higher than non-targeting control. XRCC6/Ku70 perturbation resulting in a much more subtle phenotype. Only one crRNA was effective for perturbing XRCC6/Ku70, and this perturbation resulted in an approximately 1.2-fold increase in HIV-1 infectivity.

**Figure 2.**
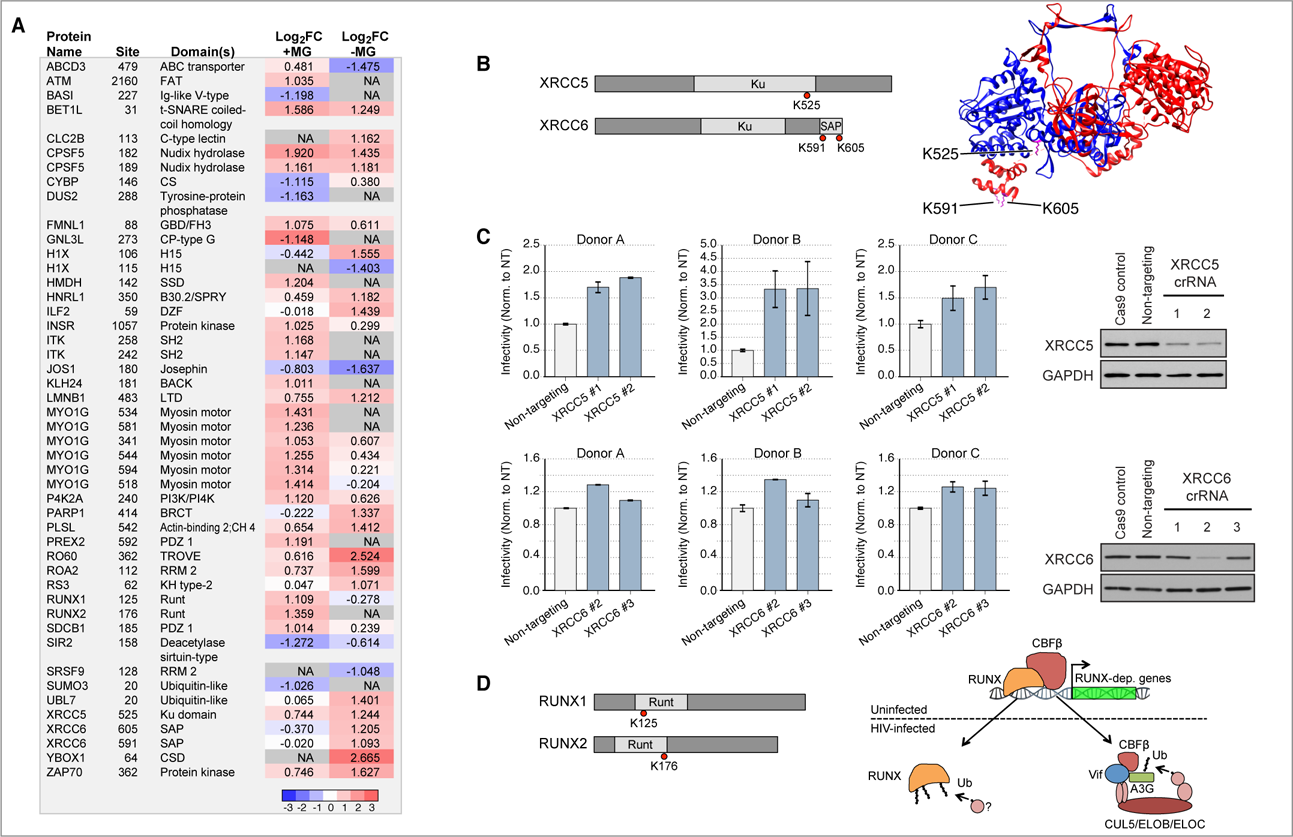
Protein domain analysis of HIV-1 regulated ubiquitination (A) Table of HIV-1-regulated ubiquitination sites occurring within defined protein domains (B) Protein domain schematic of XRCC5/Ku80 and XRCC6/Ku70 (left); structure of the XERCC5/Ku80 (blue) and XRCC6/Ku70 (red) heterodimer with regulated ubiquitination sites highlighted in magenta (PDB entry 1jeq) (Walker et al., 2001) (C) CRISPR/Cas9 editing and HIV-1 infection rates normalized to non-targeting control for XRCC5/Ku80 and XRCC6/Ku70 in human primary CD4+ T cells and Western blot analysis from a representative donor (D) Protein domain schematic and model of RUNX ubiquitination by Vif-mediated CBFβ sequestration.

We also identified 2 regulated ubiquitination sites within the Runt domains of RUNX1 and RUNX2, transcription factors that form a heterodimeric complex with core-binding factor β (CFBβ). CFBβ is hijacked by HIV-1 Vif to enhance Vif stability and aid in recruitment of the CRL5^ELOB/ELOC^ E3 ubiquitin ligase complex. These RUNX ubiquitination sites were differentially increased exclusively in the presence of proteasome inhibition (Figure 2D). RUNX1 dimerization with CFBβ protects RUNX1 from ubiquitination and degradation by the proteasome and our finding that RUNX proteins are ubiquitinated in HIV-1-infected cells in the presence of proteasome inhibitor is consistent with a model where HIV-1 Vif sequesters CFBβ away from its RUNX cofactors (Huang et al., 2001; Kim et al., 2013).

### HIV-1 Requires Aurora Kinase A for Replication in Primary CD4+ T Cells and Monocyte-Derived Macrophages

We next implemented a global phosphoproteomics approach to globally characterize changes in phosphorylation in response to HIV-1 infection (Figure 3A). Phosphoproteomics analysis was performed using label-free quantification to allow for comparisons across multiple conditions. As above for ubiquitinat remnany analyses, Jurkat T cells were infected with *Env*-deficient HIV-1 that was pseudotyped with VSVG and mock-infected with an *Env*-deficient HIV-1 that was not pseudotyped, as above for ubiquitin remnant analyses. We also infected Jurkat T cells with *Env* and *Vif*-deficient HIV-1 that was VSVG-pseudotyped to determine the impact of *Vif* deletion on the host cellular phosphoproteome. Cells were lysed 24 hours post-infection, digested with trypsin, and then subjected to Fe^3+^-immobilized metal affinity chromatography (IMAC) enrichment of phosphopeptides (Mertins et al., 2018) Enriched phosphopeptides were analyzed on an Orbitrap Fusion mass spectrometry system. Data were processed as above, first with MaxQuant and then by MSstats. This analysis identified 8,278 phosphorylation sites on 2,704 proteins, of which 366 sites were differentially increased and 406 sites were differentially decreased by at least 2-fold with an adjusted p-value < 0.05 in response to HIV-1 infection (Figure 3B, Supplementary Table S2).

**Figure 3.**
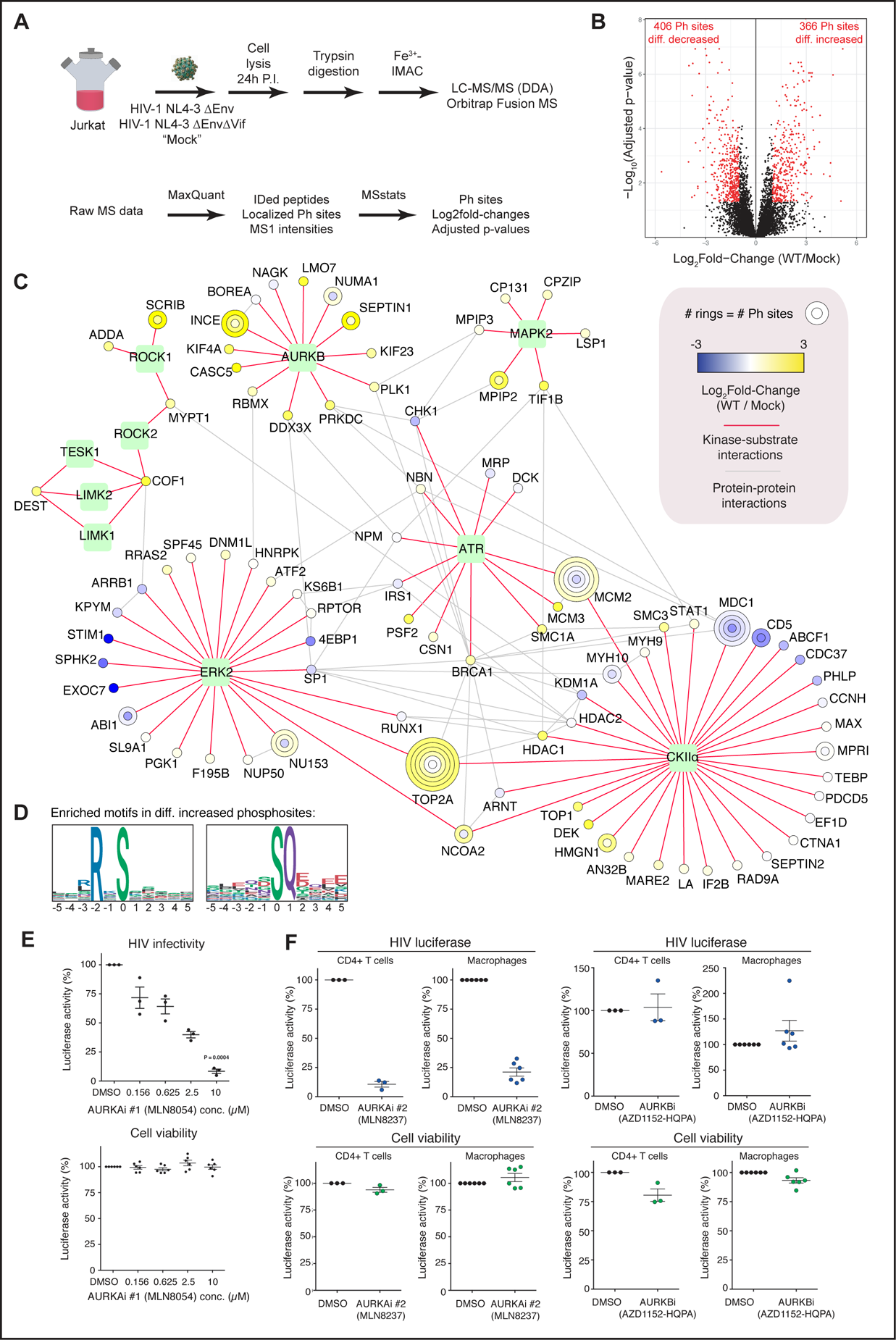
Phosphoproteomics analysis reveals a requirement for Aurora kinase A in HIV-1 infection (A) Schematic of experimental and data analysis workflows implemented (B) Volcano plot of phosphorylation changes in response to HIV-1 infection. Phosphorylation sites considered differentially increased or decreased with a |log_2_fold-change| > 1 and adjusted p-value < 0.05 are highlighted in red. (C) Kinase-substrate network for kinases with differential activities in HIV-1-infected cells (see legend for node and edge definitions). (D) Logo representations of motifs identified to be overrepresented in differentially increased phosphosites by Motif-X. (E) HIV-1 infectivity and cell viability in response to MLN8054 titration in human primary CD4+ T cells, normalized to DMSO control (F) HIV-1 infectivity and cell viability in response to MLN8237 and AZD1152-HPQ treatment in human primary CD4+ T cells and monocyte-derived macrophages, normalized to DMSO control

In order to identify kinases that may be differentially active in HIV-1 infection, phosphoproteomics data were compared to the PhosphoSitePlus kinase-substrate table curated from literature (Hornbeck et al., 2015). A hypergeometric test was employed for each kinase in the table to calculate the probability that the number of differentially increased or decreased phosphorylation sites on substrates of each kinase data was more than would be expected at random. This analysis identified 10 kinases that were regulated with a p-value < 0.05: AURKB (p = 1.04×10^-5^), ROCK2 (p = 0.0026), TESK1 (p=0.0026), LIMK1 (p=0.0026), LIMK2 (p=0.0026), ROCK1 (p = 0.014), ATR (p=0.019), MAPK2 (p=0.043), ERK2 (p=0.048), and CKIIα (p=0.049). A network view of differentially regulated kinases and their substrates is illustrated in Figure 3C. We find topoisomerase II alpha (TOP2A) phosphorylation increased upon HIV-1 infection at multiple sites, consistent with reports that have found phosphorylated TOP2A in both HIV-infected cells and in virions (Kondapi et al., 2005; Matthes et al., 1990) phosphorylation-based activation has been associated with either casein kinase II alpha (CKIIα) or extracellular signal-regulated kinase 2 (ERK2) activity, however our analysis indicates that CKIIα activity is increased while ERK2 activity is decreased by HIV-1 α infection (Ackerman et al., 1985; Shapiro et al., 1999) TOP2A and CKIIα inhibitors have both been demonstrated to inhibit HIV-1 replication at the levels of integration and transcriptional activation, respectively (Critchfield et al., 1997). Our analysis also predicted increased activity for a cluster of kinases including ROCK1, ROCK2, LIMK1, LIMK2, and TESK1, that target overlapping substrates. ROCK1 and LIMK1 have been described to modulate retrovirus particle release, while cofilin (COF1) phosphorylation, which may be targeted by many kinases, is activated by HIV envelope-CXCR4 signaling in order to overcome cortical actin restriction in resting CD4+ T cells (Wen et al., 2014; Yoder et al., 2008). Our infection conditions with VSVG-pseudotyping would bypass CXCR4 signaling, which suggests that COF1 is activated by an alternative pathway in HIV-1 infection.

Motif enrichment analysis using Motif-X identified two motifs overrepresented in differentially increased phosphorylation sites (Figure 3D): an RxS* motif and an S*Q motif. No significantly overrepresented motifs were identified in differentially decreased phosphorylation sites. RxS* motifs are common among kinase preferences and may reflect activity of Aurora kinases A and B, p21-activated kinases, AMP-activated kinase, protein kinase A, and Ca^2+^-calmodulin-dependent kinases. S*Q motifs frequently represent targets of DNA damage kinases ATM and ATR and likely reflect the activation of the DNA damage response pathway by HIV-1 Vpr (Kim et al., 1999; O’Neill et al., 2000).

Our differential kinase activity analysis identified Aurora kinase B as the most significantly regulated kinase based on its substrate phosphorylation profiles. Protein abundance analysis indicated that both Aurora kinases A and B were differentially increased in HIV-1-infected cells (Table S3). As Aurora kinases A and B (AURKA and AURKB) also have overlapping substrate preferences, we tested the impact of Aurora kinase inhibitors on HIV-1 infection in primary human CD4+ T cells. Cells were pre-treated for 1 hour with an Aurora kinase inhibitor targeting both AURKA and AURKB, MLN8054, then infected with HIV-1 containing a luciferase reporter in the Nef locus (Manfredi et al., 2007). We observed cell cycle defects with MLN8054 treatment longer than 24 hours and therefore measured luciferase activity at 24 hours and confirmed normal cell cycle progression (data not shown). MLN8054 inhibited HIV-1 infection in a dose-dependent manner with an EC50 of approximately 500 nM. We next tested selective inhibitors of AURKA and AURKB (Figure 3F). MLN8237 is 200-fold more selective for AURKA than AURKB and inhibited HIV-1 infection at 5 μM in both human primary CD4+ T cells and monocyte-derived macrophages with no effect on cell viability. AZD1152-HQPA (Barasertib), an AURKB inhibitor that is (Tomita and Mori, 2010) 3000-fold more selective for AURKB than AURKA, had no effect on HIV-1 replication but did reduce cell viability by 10-20% in human primary CD4+ T cells and monocyte-derived macrophages (Wilkinson et al., 2007). Collectively, these data uncover several substrate-kinase networks impacted during HIV-1 infection and demonstrate the essential role of AURKA signaling for productive infection.

### Global Protein Abundance Integration Identifies Ubiquitination Events Associated with Protein Degradation

To identify ubiquitination events that are associated with protein degradation in the same biological system, inputs from the ubiquitin remnant immunoaffinity enrichment procedure for experiments performed in the absence of proteasome inhibitor were subjected to hydrophilic interaction chromatography (HILIC) fractionation and mass spectrometry analysis (Figure 4A, Supplementary Table S3). Protein abundance data were processed similarly as above, first with MaxQuant to identify proteins and extract SILAC ratios, and then with MSstats to estimate log_2_fold-changes and adjusted p-values. 6,056 proteins could be quantified in sufficient biological replicates for statistical analysis. Protein abundance analysis recapitulated effects for several factors that have been reported to be degraded by HIV-1 infection, including BST2/tetherin that is degraded by Vpu, HLTF that is degraded by Vpr, and two members of the PP2A B56 variable regulatory subunit family (peptides identified could not distinguish delta and gamma family members) – that are degraded by Vif (Figure 3B) (Laguette et al., 2014; Lahouassa et al., 2016a; Lv et al., 2018; Yan et al., 2018).This analysis also recapitulated stabilization of β-catenin that has been reported to be a result of sequestration of the Cullin-1/Skp1/βTrCP ubiquitin ligase by the Vpu protein (Besnard-β Guerin et al., 2004).

**Figure 4.**
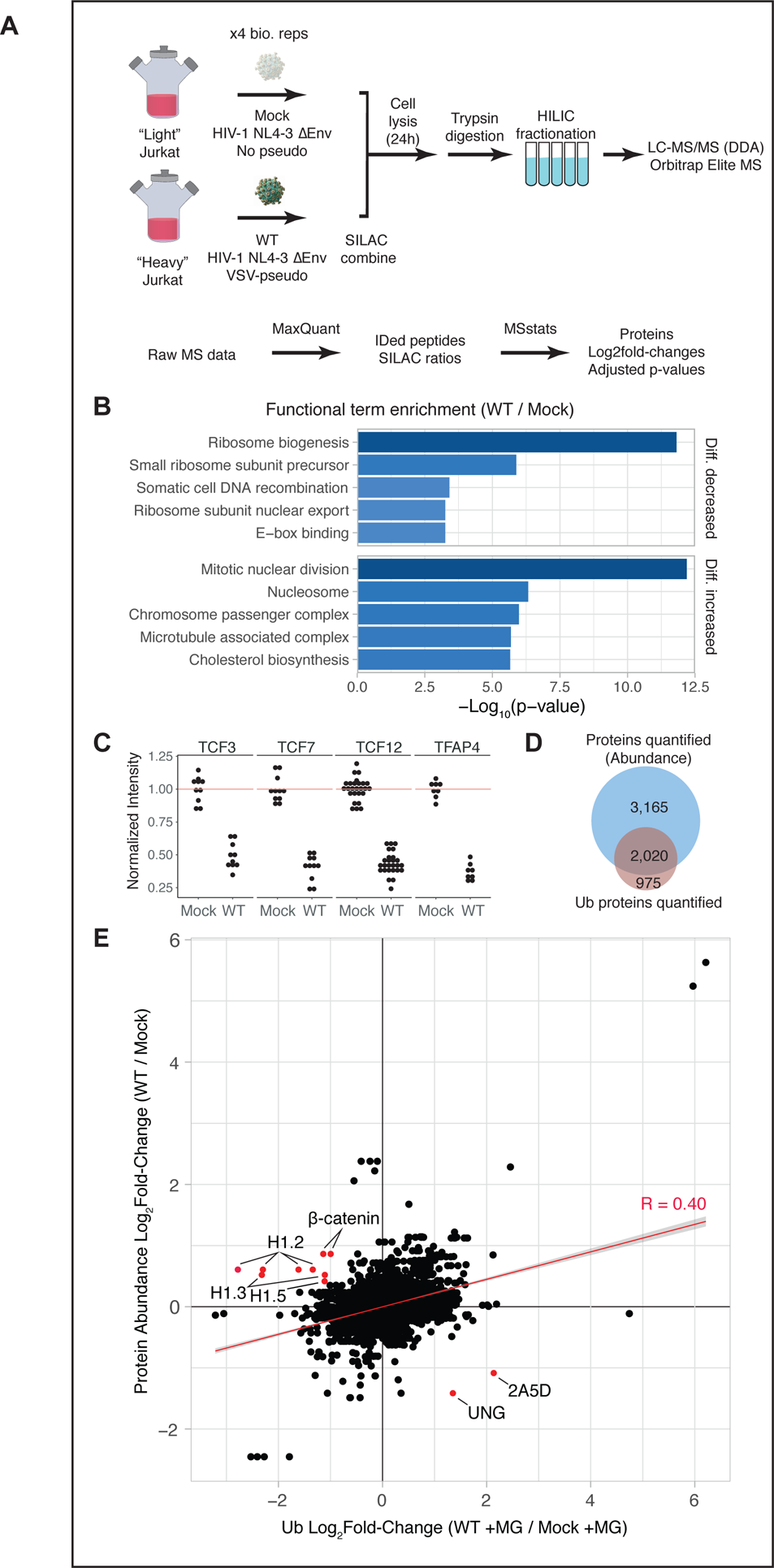
Quantification of protein abundance and integration with ubiquitination profiles (A) Schematic of experimental and data analysis workflows implemented (B) Pathway and process enrichment of differentially increased and decreased protein abundance changes (C) Dot plots of normalized peptide intensities for transcription factors downregulated by HIV-1 infection. Each dot represents an individual peptide in an singlebiological replicate. Data are normalized such that the median of uninfected intensities averages to 1 (indicated by a red line). (D) Overlap of proteins detected by ubiquitination analysis and protein abundance analysis (E) Scatterplot of changes in ubiquitination versus changes in protein abundance. Proteins with differential ubiquitination greater than 2-fold and with differential protein abundance with an unadjusted p-value < 0.05 are highlighted in red and labeled.

Pathway and process enrichment analysis by Metascape was performed for proteins with an unadjusted p-value < 0.05 regardless of their fold-change. This analysis revealed a strong enrichment for upregulation of abundance for proteins involved with mitotic processes including mitotic nuclear division (p-value = 6.41×10^-13^) and protein components of the nucleosome (HIST1H1C, HIST1H1D, HIST1H1B, H1FX, H3F3A, HIST4H4, HP1BP3, p = 4.71×10^-7^), contrasting with the decreased ubiquitination observed for many of the same histone H1 variants. Proteins that were differentially decreased in abundance were associated with several ribosome processes including ribosome biogenesis (p-value = 1.54×10^-12^), small ribosome subunit precursor (p-value = 1.30×10^-6^), and ribosome subunit nuclear export (p-value = 5.51×10^-4^). Differentially decreased proteins were also associated with E-box binding (PPARG, TCF3, TCF12, TFAP4, p = 5.51×10^-4^) and highlights several transcription factors downstream of β-catenin signaling that were found to be differentially decreased by HIV-1 infection: TFAP4, TFE3/TCF3, TCF7, and HTF4/TCF12 were all differentially decreased in abundance (Figure 4C).

Of the 2,995 proteins for which we quantified ubiquitination changes, 2,020 (67%) were also quantified by protein abundance analysis (Figure 4D). We sought to integrate ubiquitination and protein abundance data in order to identify proteins whose abundance may be regulated by ubiquitination and degradation by the proteasome. To that end, we extracted proteins with ubiquitination sites that changed greater than 2-fold in the presence of proteasome inhibition where their protein abundance, performed in the absence of proteasome inhibition, changed in the opposite direction with an unadjusted p-value < 0.05 (Figure 4E). This analysis highlighted several proteins regulated by HIV-1-mediated ubiquitination and degradation, including β-catenin, PP2A-B56 (2A5D), and UNG, which is targeted for ubiquitination and degradation by Vpr (Besnard-Guerin et al., 2004; Bouhamdan et al., 1996; Greenwood et al., 2016; Selig et al., 1997. Additionally, this analysis identified histone H1 variants H1.2, H1.3, and H1.5 as being stabilized by an apparent downregulation of their ubiquitination.

### HIV-1 Accessory Genes Contribute to a Network of Ubiquitination

We next sought to map ubiquitination sites to specific HIV-1 accessory genes by comparing cells infected with wild-type HIV-1 to HIV-1 mutants that were deficient in expression of *Vif*, *Vpu*, or *Vpr* (Figure 5A, Supplementary Table S4). Experiments with accessory gene-deficient HIV-1 mutant viruses were performed as above for wild-type vs. mock infection experiments using SILAC for quantification, with 4 biological replicates of each wild-type vs. mutant HIV-1 comparison in the presence and absence of proteasome inhibitor (MG-132). Data were processed with MaxQuant and MSstats as described above. This analysis quantified 5,446, 8,545, and 7,729 ubiquitination sites in response to wild-type vs. *Vif*-, *Vpr*-, and *Vpu*-deficient HIV-1, respectively, with 9,869 ubiquitination sites quantified in total (Figure 5B). Using the same criteria for differentially abundant ubiquitination in the wild-type vs. mock comparison (i.e., |log_2_fold-change| > 1.0 and adjusted p-value < 0.05) we identified ubiquitination sites that were regulated by deletion of *Vif*, *Vpu*, or *Vpr* that were similarly regulated comparing wild-type to mock infection. These changes are represented in a network view in Figure 5C overlaid with multivalidated protein-protein interactions from BioGRID (Oughtred et al., 2018) analysis recapitulates the previously reported Vif-dependent ubiquitination and degradation of PP2A-B56 family members delta and gamma (2A5D and 2A5G), as well as the PP2A catalytic subunit (2AAA) (Greenwood et al., 2016). Some ubiquitination sites appeared to be differentially abundant in response to deletion of multiple accessory genes. Two proteins in the cholesterol biosynthesis pathway were differentially abundant in response to all three accessory gene deletions including one site on squalene synthase (FDFT) and two sites on the cytoplasmic hydroymethylglutaryl-CoA synthase (HMCS1). Two ubiquitination sites on RUNX1 were observed to be regulated by *Vpu* deletion, with one of those sites similarly regulated by *Vif* deletion. Ubiquitination sites on protein components of both the 40S and 60S ribosome subunits were regulated by all three accessory genes in both directions. Vif-regulated ubiquitination sites on ribosomal protein S3, S16, L8, and L36 were observed to increase in HIV-1 infection, while 4 of 5 Vpr-regulated ubiquitination sites on ribosomal proteins were downregulated. Finally, we mapped the activity of histone H1 ubiquitination on variants H1.2 and H1.4 specifically to the *Vpr* gene.

**Figure 5.**
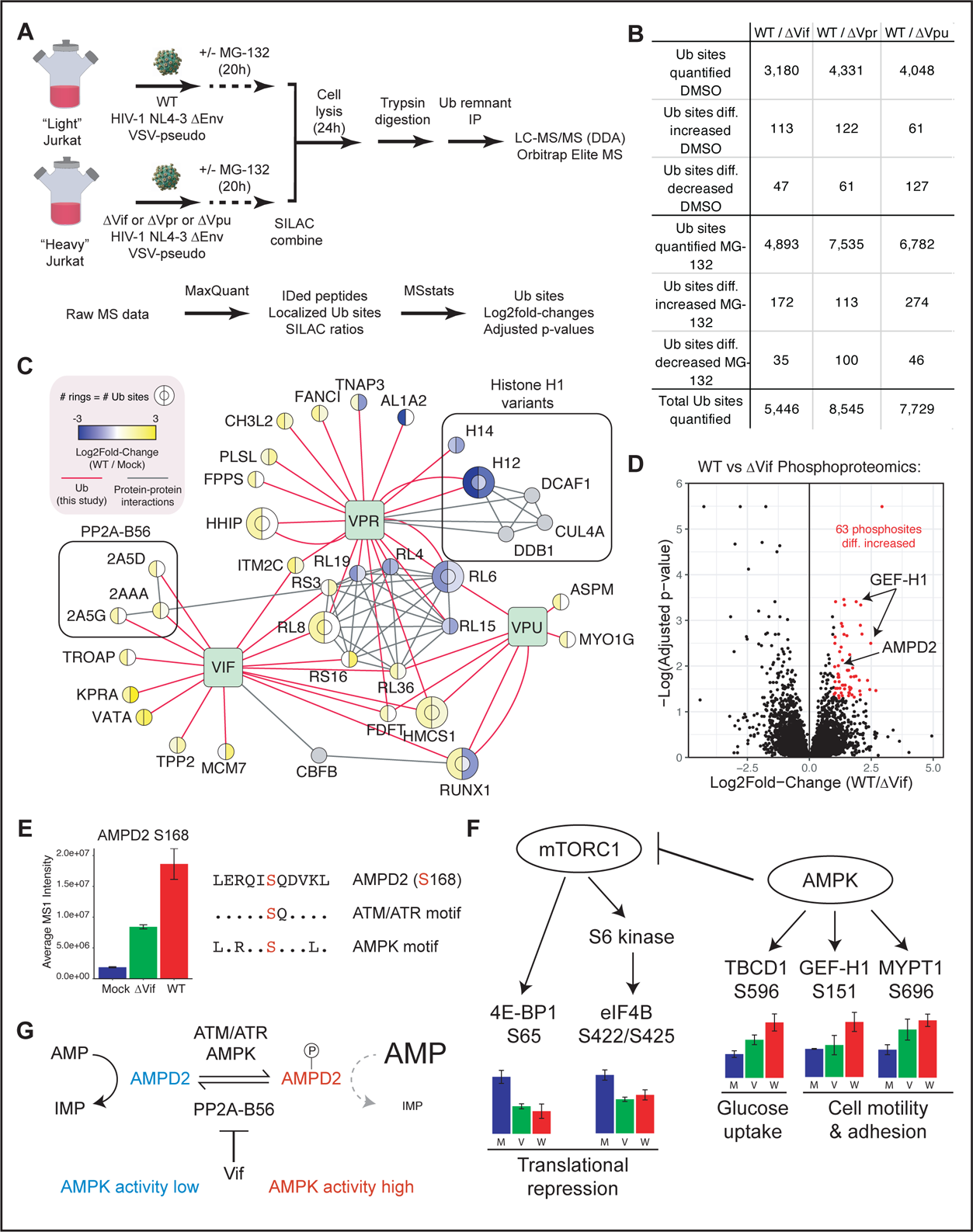
HIV-1 Vif targets PP2A-B56 for ubiquitination and results in hyperphosphorylation of AMPD2 (A) Schematic of experimental and data analysis workflows implemented (B) Summary of ubiquitination site identification and differentially increased and decreased ubiquitination sites by mutant viruses (C) Network view of putative Vif-, Vpr-, and Vpu-dependent ubiquitination. See legend for node and edge definitions. (D) Volcano plot of phosphorylation changes comparing cells infected with wild-type and *Vif*-deficient HIV-1. Phosphosites that were differentially increased in cells infected with wild-type HIV-1 (i.e., log_2_fold-change WT/Δ highlighted in red. Phosphorylated proteins containing the PP2A-B56 LxxIxE binding motif are labeled and named. (E) Bar plots of AMPD2 S186 phosphorylation intensities after normalization by MSstats (left); sequence region of AMPD2 S186 and kinase consensus motifs (right). (F) Schematic of mTORC1 and AMPK pathways. Bar plots indicate phosphorylation site intensities after normalization by MSstats; M=mock, V=*Vif*-deficient HIV-1-infected, W-wild-type HIV-1-infected; error bars indicate standard deviation. (G) Model representing how AMPD2 phosphorylation is modulated by HIV-1 and impacts AMPK activity.

### PP2A-B56 Degradation by Vif Enhances AMPK Signaling by Modulating AMPD2 Phosphorylation

To identify phosphorylation sites that may be regulated by Vif-mediated degradation of PP2A-B56 we compared phosphoproteomics profiles of cells infected with wild-type and *Vif*-deficient HIV-1 using label-free proteomics. This analysis identified 63 phosphorylation sites on 53 proteins that were at least 2-fold more abundant in cells infected with wild-type compared to *Vif*-deficient HIV-1 with an adjusted p-value < 0.05 (Figure 5D). PP2A-B56 contains a conserved, surface-exposed pocket that binds to proteins containing a LxxIxE consensus motif (Hertz et al., 2016). Of the proteins whose phosphorylation status was affected by *Vif* deletion two proteins contained LxxIxE motifs: 1) the Rho guanine exchange factor GEF-H1, which has been validated as a PP2A-B56 substrate, and 2) AMP deaminase 2 (AMPD2) (Hertz et al., 2016). AMP deaminase catalyzes the deamination of AMP to IMP. AMP deaminase inhibition leads to an increase in AMP, an increased AMP:ATP ratio, and increased activation of AMP-activated protein kinase, a critical regulator of cellular energy homeostasis (Plaideau et al., 2014). We found several substrates of AMPK that were phosphorylated in a manner consistent with AMPK activation including TBCD1, GEF-H1, and MYPT1 (Figure 5F).

Furthermore, AMPK activity is associated with inhibition of mTOR signaling, and we found that two classical mTORC1 substrates, 4E-BP1 and eIF4B, were less phosphorylated in conditions with increased AMPK activity. Sequence analysis of the AMPD2 phosphorylation site at position S168 revealed overlapping motifs for both ATM/ATR kinases (S*Q) and AMPK (LxRxxS*xxxL) (Figure 5E). We propose a model whereby AMPD2 is driven towards a hyperphosphorylated state initially by ATM and/or ATR kinases, which are activated by Vpr-induced activation of the DNA damage response pathway (Figure 5G). PP2A-B56 reverses AMPD2 phosphorylation, while Vif degrades PP2A-B56 to retain AMPD2 phosphorylation. Phosphorylated and inactive AMPD2 leads to increased levels of cellular AMP, thus activating AMPK in a manner that upregulates processes that provide energy stores for HIV-1 replication.

### Vpr Inhibits Ubiquitination of Histone H1 Variants by Inhibiting CRL4^DCAF1^

Histone H1 variants were found to be heavily modified by PTMs that were regulated by HIV-1 infection. Figure 6A illustrates the phosphorylation and ubiquitination sites identified on all histone H1 variants and their differential abundance in cells infected with HIV-1 compared to mock virus. The central globular domain is highly conserved among histone H1 variants and many modified peptides in this region could not distinguish precisely which variants were modified. The N- and C-terminal regions, however, are more variable and allowed us to identify increased phosphorylation and decreased ubiquitination of histone H1 variants H1.2, H1.3, H1.4, and H1.5 at positions K17 and K21 in response to HIV-1 infection. Similarly, phosphorylation at position S2 was increased in histone H1 variants H1.2, H1.2, H1.4, and H1.5. No modifications were observed on histone H1 variants H1.0 or H1.1. Histone H1.1 was not detected in protein abundance analysis and histone H1.0 was detected with far fewer peptides than the somatic variants, so it is possible that these histone H1 isoforms may be modified but are below the detection limit of our assay.

**Figure 6.**
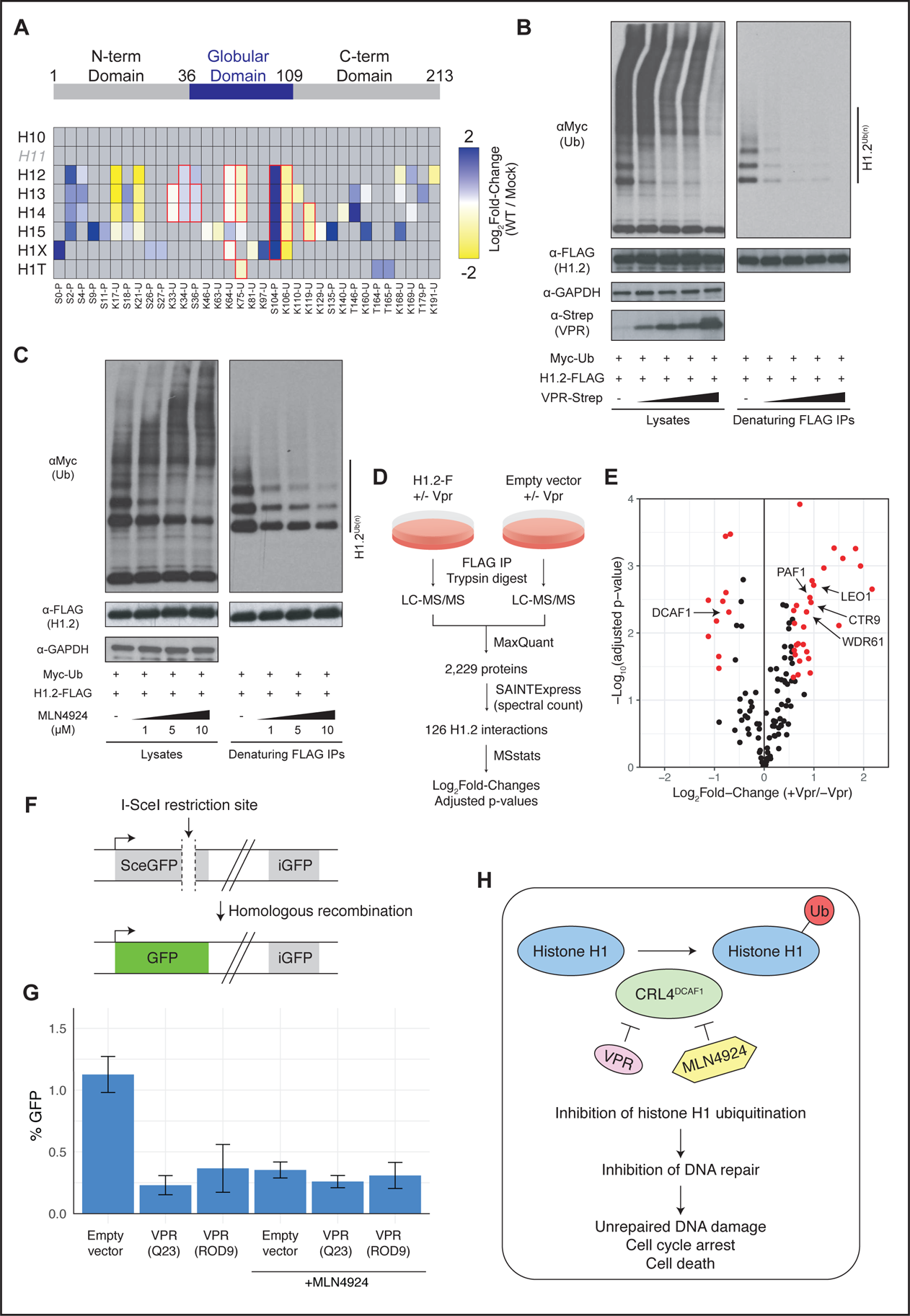
Vpr inhibits ubiquitination of histone H1 variants by CRL4^DCAF1^ (A) Schematic of histone h1 protein domains (top; positions indicated are for histone H1.2). Heatmap of phosphorylation (P) and ubiquitination (U) changes on histone H1 variants. All sites are aligned to the sequence positions of histone H1.2. Cells with a red outline were identified by peptides that cannot distinguish between histone variants. (B) Denaturing ubiquitin immunoprecipitation analysis of histone H1.2 with Vpr titration (C) Denaturing ubiquitin immunoprecipitation analysis of histone H1.2 with MLN4924 titration (D) Schematic of experimental and data analysis workflows for quantitative affinity purification and mass spectrometry analysis of histone H1.2 in the presence and absence of Vpr co-expression (E) Volcano plot of histone H1.2 protein binding changes in response to Vpr expression. Proteins with |log_2_fold-change| > 0.58 (i.e., 1.5-fold change) and adjusted p-value < 0.05 are highlighted in red. (F) Schematic of a fluorescence-based assay for homology-directed repair of DNA double strand breaks. (G) Homology-directed repair assay of cells in response to VPR expression and MLN4924 treatment. Bar heights represent an average of three biological replicates and error bars indicate the standard deviation. (H) A model for HIV-1 VPR-mediated inhibition of histone H1 ubiquitination, DNA repair, cell cycle arrest, and cell death.

Our analysis of cells infected with *Vif*-, *Vpu*-, and *Vpr*-deficient HIV-1 revealed that ubiquitination of histone H1 variants H1.2 and H1.4 is Vpr dependent. We validated that Vpr expression reduced histone H1 ubiquitination by performed denaturing immunoprecipitation of FLAG-tagged histone H1.2 that was co-transfected with Myc-tagged ubiquitin and increasing amounts of Vpr (Figure 6B). Histone H1 variants physically interact with the CRL4^DCAF1^ ubiquitin ligase, which suggests two likely scenarios: 1) CRL4^DCAF1^ is a histone H1 ubiquitin ligase whose activity towards histone H1 is reduced by Vpr, or 2) Vpr hijacks CRL4^DCAF1^ to inhibit a histone H1 ubiquitin ligase (Huang et al., 2001; Kim et al., 2013a). To discern which of these scenarios occurs we used a NEDD8 activating enzyme inhibitor, MLN4924, which inactivates Culling RING-type E3 ubiquitin ligases by preventing their NEDDylation-based activation (Soucy et al., 2009). MLN4924 treatment reduced histone H1.2 ubiquitination in a dose-dependent manner similar to Vpr expression. This finding is consistent with the first scenario described above whereby CRL4^DCAF1^ is a histone H1 ubiquitin ligase that is inhibited by Vpr (Figure 6C).

To understand how Vpr inhibits CRL4^DCAF1^ ubiquitination of histone H1 variants we next employed quantitative affinity purification and mass spectrometry (AP-MS) analysis of histone H1.2 to quantify changes in histone H1.2 protein interactions in response to Vpr expression (Figure 6D). 293T cells were transfected with FLAG-tagged histone H1.2 or an empty vector were co-transfected with Vpr or empty vector in biological triplicate for each condition. Cells were lysed and subjected to native FLAG affinity purification and mass spectrometry analysis of the eluates on an Orbitrap Fusion mass spectrometry system. Raw MS data were analyzed with a similar workflow as PTM analyses, first with MaxQuant to identify peptides and extract MS1 peak areas, then with MSstats to fit a model of all levels of variation and to test the significance of observed changes relative to the model. Additionally, we applied SAINTexpress with the spectral counting method to filter out non-specific protein interactions using the empty vector transfected cells as negative controls (Teo et al., 2014). This analysis identified 126 histone H1.2 interacting protein of which 28 and 10 were differentially increased or decreased by Vpr co-expression, respectively (Figure 6E, Supplementary Table S5). DCAF1 binding was reduced by 1.6-fold when Vpr was co-expressed, as well as three subunits of the mitochondrial ribosome. Increased binding was observed between histone H1.2 and members of the PAF1 complex (PAF1, CTR9, LEO1, and WDR61) when Vpr was co-expressed.

Histone H1 ubiquitination plays a key role in the DNA damage response by facilitating the recruitment of repair factors BRCA1 and 53BP1 to sites of damage to initiate repair processes (Thorslund et al., 2015). Vpr has been recently demonstrated to inhibit DNA repair (personal communication with Oliver Fregoso; manuscript in revision). To test whether histone H1 deubiquitination affects DNA repair processes we implemented a fluorescence-based assay of homology-directed repair (HDR) of DNA double-strand breaks (Figure 6F) (Pierce et al., 1999). In this system, I-SceI expression induces a double-strand break within a defective GFP locus that is made functional when repaired from a second GFP locus in the genome. Neither GFP locus is functional unless HDR occurs. We found that expression of VPR derived from HIV-1 (Q23 isolate) and HIV-2 (ROD9 isolate) inhibited HDR by up to four-fold (Figure 6G). A similar reduction in HDR was observed when cells were treated with MLN4924, indicating that Cullin inhibition – and not Cullin-mediated degradation of a target protein – causes HDR impairment (Figure 6G).

## DISCUSSION

In this study, we present a quantitative analysis of ubiquitination, phosphorylation, and protein abundance changes in response to HIV-1 infection. We demonstrate strategies to integrate and interpret complex proteomics data in order to identify host cellular pathways that are targeted by HIV-1 and to determine the molecular mechanisms by which HIV-1 perturbs their function. Our findings also provide insights regarding host cellular pathways that are independent of HIV-1 infection.

### Determining PTM Enzyme-Substrate Relationships by Shotgun Proteomics

We demonstrate the utility of global PTM analyses to identify PTM enzyme-substrate relationships. Physical interactions between PTM enzymes and their substrates are often low affinity and/or transient, thus approaches to identify these relationships by affinity purification or immunoprecipitation approaches are often unsuccessful. By perturbing PTM enzymes or the HIV-1 proteins that hijack them and globally profiling the cognate PTM type, we identified PP2A-B56 as a ubiquitination substrate of Vif-CRL5^ELOB/ELOC^, AMPD2 as a phosphorylation substrate of PP2A-B56, and histone H1 somatic variants as ubiquitination substrates of CRL4^DCAF1^ that are inhibited by Vpr. These examples demonstrate that current proteomics technologies are sufficient to map functional PTM enzyme-substrate relationships, setting the stage for more systematic, comprehensive studies of these relationships.

### Integrating ubiquitination profiles with proteasome perturbations and protein abundance measurements

Our ubiquitination analysis identified many proteins that were ubiquitinated on multiple resides and we demonstrated that a vast majority (>75%) of those sites respond similarly to HIV-1 infection. This has implications on the experimental approaches intended to test the biological function of individual ubiquitination sites: if the exact site of ubiquitination is flexible then mutating residues found to be ubiquitinated is unlikely to elicit an effect in a functional assay. Along this line of reasoning, we cannot conclude that the differentially ubiquitinated sites we identified on XRCC5/Ku80 in HIV-1-infected cells have a functional impact on its restriction activity. Rather, our discovery that the Ku complex is ubiquitinated in HIV-1-infected cells led us to ask two questions: 1) whether Ku possesses restriction activity against HIV-1, and 2) whether that activity may be counteracted by ubiquitination. We addressed the first question by perturbing Ku in a CRIPSR/Cas9 system in primary CD4+ T cells and demonstrating that it behaves like a restriction factor. To properly answer the second question it will be of interest to identify the upstream ubiquitin ligase and perturb its function to measure the effects HIV-1 infectivity.

We have previously demonstrated that differential ubiquitination of APOBEC3C and CD4 in HIV-1-infected cells could only be observed in the presence of proteasome (Ball et al., 2016. By combining ubiquitination profiles of proteasome-inhibited, HIV-1-infected cells with protein abundance profiles of HIV-1-infected cells in which the proteasome was not inhibited, we rapidly identified PP2A-B56, UNG, β-catenin, and histone H1 isoforms as ubiquitination events that are destined for degradation by the proteasome. While proteasome inhibition is essential for identifying ubiquitination substrates that are targeted for proteasomal degradation, we found a low overlap between ubiquitination changes in the presence and absence of proteasome inhibition, suggesting that proteasome inhibition has profound effects on cellular ubiquitination profiles. This presents a challenge when interpreting ubiquitination profiles for proteins such as RUNX1 and RUNX2, where differential ubiquitination is observed only the presence of proteasome inhibition but for which there are no observable changes in protein abundance. One possible explanation could be that only a subpopulation of these proteins is targeted for ubiquitination and degradation such that the total steady-state abundance of these proteins is not significantly perturbed.

In the case of RUNX proteins, HIV-1 Vif recruits the RUNX cofactor CFBβ into a complex with the CUL5^EloB/EloC^ ubiqiuitin ligase in order to target APOBEC3 proteins for proteasomal degradation (Jäger et al., 2012). CFBβ protects RUNX proteins from ubiquitination and degradation and our findings are consistent with a model where Vif sequesters CFBβ and exposes RUNX proteins to ubiquitination (Huang et al., 2001; Kim et al., 2013a). That RUNX proteins are not observed to be differentially abundant in HIV1-infected cells could imply that Vif is targeting a specific CBFβ-RUNX subpopulation for disruption. It will be of interest to identify the characteristics of that population (e.g., genomic location, post-translational modifications, or relevant cofactors) and a mechanism for how a specific population is targeted by Vif.

### Phosphoproteomics analysis identifies a requirement for Aurora kinase A activity in HIV-1 infection

Phosphoproteomics analysis and subsequent kinase-substrate analysis revealed a strong enrichment for Aurora kinase B substrates in HIV-1-infected cells. However, knowledge of kinase-substrate relationships is heavily biased towards well-studied kinases in contexts that have been the subject of systems-wide interrogation. In fact, we found the protein abundances of both Aurora kinases A and B are differentially increased in HIV-1-infected cells, but only Aurora kinase B was identified from our kinase-substrate analysis. Indeed, inhibition of Aurora kinase B appears to have no effect on HIV-1 replication, but inhibition of Aurora kinase A, which shares some substrate specificity with Aurora kinase B, profoundly inhibits infection. These findings highlight a gap in knowledge regarding Aurora kinase A substrates and highlights the limitations of existing empirical resources regarding kinase-substrate relationships.

Interestingly, Aurora kinases A and B have both been reported to be inhibited by the DNA damage response (DDR) pathway, while it is well established that the HIV-1 Vpr protein activates DDR (Krystyniak et al., 2006; Monaco et al., 2005). How the Aurora kinases maintain their activity in the presence of activated DDR also remains to be determined.

### Vif Modulates Cellular Energy Homeostasis via PP2A-B56 Degradation

AMPK is a master regulator of cellular energy homeostasis that is conserved throughout the three domains of life (Hardie, 2007; Roustan et al., 2016). Because it is activated by AMP and inhibited by ATP, AMPK becomes activated under conditions where energy is depleted and functions to restore energy levels by inhibiting energy-consuming processes (e.g., protein synthesis and fatty acid synthesis) and activating energy-producing processes (e.g., glucose uptake and glycolysis). AMP deaminases control cellular levels of AMP, which are in turn sensed by AMPK. AMPK and AMP demainases have been demonstrated to counteract each other; silencing of AMPK activates AMP deaminase activity and vice versa (Lanaspa et al., 2012; Plaideau et al., 2014). Genotoxic stressors including etoposide, doxorubicin, and ionizing radiation also activate AMPK (Fu et al., 2008; Ji et al., 2010; Sanli et al., 2010)

Our phosphoproteomics analysis identified an AMPD2 phosphorylation site at S186 that was differentially increased by HIV-1 infection and enhanced by Vif. AMPD2 contains a PP2A-B56 binding motif, thus it is likely a direct target of PP2A-B56 activity that is hyperphosphorylated when PP2A-B56 is degraded by Vif. We find evidence of AMPK activation in the phosphorylation of its downstream substrates, as well as evidence of AMPK-mediated mTORC1 inhibition demonstrated by inhibition of mTORC1 substrates. These findings suggest that AMPD2 phosphorylation inhibits its activity but further experiments will be required to confirm this mechanism and to determine if S186 is the only phosphorylation site on AMPD2 that regulates its activity. The AMPD2 S186 phosphorylation site contains signatures of both ATM/ATR and AMPK consensus motifs and could provide a mechanism for AMPK activation under conditions of genotoxic stress. It is well established that the HIV-1 Vpr activates the DDR pathway rapidly upon infection of cells. Thus, HIV-1 induction of DDR may provide an initial activation phosphorylation of AMPD2 and concomitant activation of AMPK signaling, followed later by Vif-mediated degradation of PP2A-B56, increased AMPD2 phosphorylation, and further upregulation of AMPK activity during post-integration steps of infection.

AMPK activation by viruses provides a rapid mechanism for simultaneously disabling host protein synthesis and increasing energy stores available for energy-intensive viral production. However, AMPK-mediated inhibition of certain biosynthetic processes would be expected to have a negative impact on viral replication. Of note, we did not observe inhibitory phosphorylation of the canonical AMPK substrate acetyl-CoA carboxylase (ACC), which catalyzes the rate-limiting step for fatty acid. This may suggest that HIV-1 has evolved separate mechanisms for maintaining biosynthetic activities in the presence of active AMPK. Intriguingly, AMPD2 deficiency in patients is associated with early-onset neurodegeneration that was found to be associated with defective GTP biosynthesis (Akizu et al., 2013. GTP is an important activator of the HIV-1 restriction factor SAMHD1, thus GTP depletion via AMPD2 inactivation may impair its capacity to restrict HIV-1 (Amie et al., 2013)

### Vpr Modulates DNA Damage Signaling by Inhibiting the Activity of CRL4^DCAF1^ Towards Histone H1 Somatic Variants

Histone H1 ubiquitination plays a key role in the DNA damage response by facilitating the recruitment of repair factors BRCA1 and 53BP1 to sites of damage to initiate repair processes (Thorslund et al., 2015). We found that Vpr inhibits CRL4^DCAF1^-mediated histone H1 ubiquitination and homology-directed repair (HDR) of double-strand DNA breaks. This is consistent with reports that HIV-1-infected cells accumulate double-strand DNA breaks in a Vpr-dependent manner and that even latently infected cells exhibit defects in DNA repair (Piekna-Przybylska et al., 2017; Tachiwana et al.,. Importantly, by demonstrating that MLN4924, a NEDD8 activating enzyme 1. 2006) inhibitor, phenocopies Vpr in its capacity to inhibit HDR, we propose that Vpr’s capacity to *inhibit* a Cullin activity, and not to *hijack* Cullin activity, is responsible for the HDR phenotype.

It remains to be determined how inhibition of histone H1 ubiquitination provides a benefit to HIV-1 infection. We found that histone H1.2 binding to the PAF1 complex is enhanced by Vpr. Several viruses target the PAF1 complex to suppress interferon-stimulated gene expression. Flaviviruses inhibit the recruitment of PAF1 complexes to interferon-stimulated gene locations via the NS5 protein, and the influenza virus NS1 protein mimics histones to bind PAF1 complexes and suppress antiviral gene expression (Marazzi et al., 2012; Shah et al., 2018. Therefore, it is plausible that HIV-1 perturbs PAF1 in a manner that suppresses interferon-stimulated gene expression. Histone H1 variants play important roles in chromatin accessibility, condensation, and gene regulation. It will be important to identify the whether specific genome locations in host cells are being targeted by Vpr-mediated inhibition of histone H1 ubiquitination and what the consequences of this inhibition are on gene expression profiles.

## ACKNOWLEDGEMENTS

A.M. holds a Career Award for Medical Scientists from the Burroughs Wellcome Fund, is an investigator at the Chan Zuckerberg Biohub and has received funding from the Innovative Genomics Institute (IGI) and the Parker Institute for Cancer Immunotherapy (PICI).

## AUTHOR CONTRIBUTIONS

Conceptualization, J.R.J. and N.J.K; Methodology, J.R.J., D.C.C., J.F.H, D.L., O.I.F.; Investigation, J.R.J., D.C.C., J.F.H., J.M., J.S., R.H., T.L.J., B.W.N., D.L., O.I.F.; Software, J.R.J., E.V., P.B.; Formal Analysis, J.R.J., E.V.; Writing – Original Draft, J.R.J., Writing – Review and Editing, J.R.J., N.J.K.; Visualization, J.R.J., M.S., Supervision; J.R.J., A.M., A.D.F., J.A.T.Y., O.I.F., N.J.K; Funding Acquisition, J.R.J., A.D.F., J.A.T.Y., N.J.K.

## DECLARATION OF INTERESTS

A.M. is a co-founder of Arsenal Biosciences and Spotlight Therapeutics, serves as on the scientific advisory board of PACT Pharma and is an advisor to Trizell. The Marson Laboratory has received sponsored research support from Juno Therapeutics, Epinomics, Sanofi and a gift from Gilead.

### LEAD CONTACT AND MATERIALS AVAILABILITY

Further information and requests for resources and reagents should be directed to and will be fulfilled by the Lead Contact, Jeffrey Johnson. Proviral pNL4-3 plasmids described in this paper (ΔEnv pNL4-3, ΔEnv ΔVif pNL4-3, ΔEnv ΔVpr pNL4-3, and ΔEnv ΔVpu pNL4-3) will be submitted to the NIH AIDS Reagent Program for distribution upon publication of this manuscript.

### EXPERIMENTAL MODELS AND SUBJECT DETAILS

#### Cell Lines

The Jurkat E6.1 T cell line was used as a model for T cells infected with HIV-1. Cells were grown in RPMI-1640 medium (Corning) supplemented with penicillin and streptomycin antibiotics (Gibco) and 10% fetal bovine serum (FCS, Gibco).

#### Viruses

For global proteomics analyses, the pNL4-3 plasmid (obtained form the NIH AIDS Reagent Program) of HIV-1 was modified to prevent expression of the Env glycoprotein by mutating the start codon and introducing two stop codons (Adachi et al., 1986) mutations maintain identical protein coding sequence for the overlapping *Vpu* open reading frame. *Vif*, *Vpr*, and *Vpu* were similarly mutated to disrupt their start codons and to introduce premature stop codons. For testing the effects of kinase inhibitors on HIV-1 infection in primary CD4+ T cells, cells were infected with a replication competent NL4-3/Nef-IRES/Renilla luciferase HIV-1 reporter virus that expresses Renilla lucieferase from an internal ribosome entry site, or IRES, that was provided by Sumit Chanda. For CRIPSR/Cas9 gene editing experiments in primary human CD4+ T cells, cells were infected with replication competent HIV-1 NL4-3 NEF-IRES-GFP reporter virus (obtained from the NIH AIDS Reagent Program) (Schindler et al., 2003)

## METHOD DETAILS

### Virus production

HEK293T cells were transfected in T-175 flask format with 22.02 μg of ΔEnv pNL4-3 HIV-1 provirus and 2.98 μg pcDNA/VSVg using acidified PEI pH 4 in lactate-buffered saline (LBS), which yielded a stoichiometric ratio of 3:1 provirus:envelope. Viral supernatant was collected 48 hours after transfection, cleared by centrifugation, and filtered through a 0.45 μm filter. Virus was precipitated by addition of 50% PEG-6000 and 4M NaCl to final concentrations of 8.5% and 0.3M, respectively, followed by incubation at 4 degrees for 2 hours, centrifugation and resuspension in phosphate buffer saline (PBS). Viral titer was quantified by titration on Jurkat T cells followed by fixation, staining with anti-HIV-1 Core Antigen Clone KC57 FITC (Beckman Coulter) and detection of HIV-1-infected cells by flow cytometry. *Vif*-, *Vpr*-, and *Vpu*-deficient viruses were produced in the same manner.

### HIV-1 infection of Jurkat T cells

For ubiquitination and protein abundance analyses, Jurkat cells were first labeled with light and heavy SILAC medium for two weeks. SILAC medium was comprised of RPMI-1640 lacking L-Lysine and L-Arginine supplemented with 10% dialyzed FCS, antibiotics, and natural L-Lysine and L-Arginine (for light medium) and ^13^C_6_-Lysine and ^13^C_6_,^15^N_4_-Argining for heavy medium. The efficiency of stable isotope-labeled amino acids incorporation was assessed by analyzing an aliquot of a lysate from cells grown in heavy medium to ensure that the amount of unlabeled proteins was < 1%. Infections were performed by spinoculation for 2 hours at 1400 x *g* with VSVg pseudotyped HIV-1 at MOI of 5 in the presence of 1 μg/ml PEI (Polysciences) and 16 μg/ml Polybrene (Sigma-Aldrich).

### Preparation of samples for global ubiquitination, phosphorylation, and protein abundance analysis

Infected cells were lysed in a buffer containing 8M urea, 50 mM ammonium bicarbonate, 150 mM NaCl, and PhosStop and Complete-EDTA free phosphatase and protease inhibitors (Roche) and sonicated to sheer membranes and DNA. Protein concentrations were measured by a Bradford assay. For SILAC analyses, light and heavy cells for each comparison (i.e., ΔEnv vs. mock, ⊗Env vs. ⊗Env⊗Vif, ⊗Env vs. ⊗Env⊗Vpr, ⊗Env vs. ⊗Env⊗Vpu) were combined at equal protein concentrations.

Lysates were reduced by the addition of 4 mM TCEP (Sigma) for 30 minutes at room temperature, disulfide bonds were alkylated with 10 mM iodoacetamide (Sigma) for 30 minutes in the dark at room temperature, and excess iodoacetamide was quenched with 20 mM DTT (Sigma). Lysates were diluted 1:4 in 50 mM ammonium bicarbonate and trypsin was added at a 1:100 enzyme:substrate ratio. Lysates were digested for 18 hours at room temperature with rotation. Digested lysates were acidifed with 0.1% TFA and peptides concentrated on Sep-Pak C18 solid phase extraction columns (Waters). For ubiquitin remnant analysis, an amount of digested lysate equivalent to 10 mg of protein was subjected to ubiquitin remnant immunoprecipitation according to the manufacturer’s protocol (Cell Signaling Technologies). For phosphorylation analysis, an amount of digested lysate equivalent to 1 mg of protein was lyophilized and then resuspended in a buffer containing 75% ACN with 0.1% TFA. Peptides were incubated with Fe^3+^-immobilized metal affinity chromatography (IMAC) beads, washed with the same resuspension buffer, and then phosphopeptides were eluted with 500 mM HK_2_PO_4_. For both ubiquitin remnant-enriched and phosphopeptide-enriched samples, the purified material was desalted using homemade C18 STAGE tips, evaporated to dryness, and then resuspended in 0.1% formic acid for mass spectrometry analysis (Rappsilber et al., 2007. For protein abundance analysis, an amount of digested lysate equivalent to 100 μg of protein was fractionated by hydrophilic interaction chromatography (HILIC). Peptides were injected onto a TKSgel amide-80 column (Tosoh biosciences, 2.0 mm x 15 cm packed with 5 μm particles), equilibrated with 10% HILIC Buffer A (2% ACN, 0.1% TFA) and 90% HILIC Buffer B (98% ACN, 0.1% TFA) using an AKTA P10 purifier system. The samples were then separated by a one-hour gradient from 90% HILIC Buffer B to 55% HILIC Buffer B at a flow rate of 0.3 ml/min. Fractions were collected every 1.5 min and combined into 12 fractions based on the 280 nm absorbance chromatogram. Fractions were evaporated to dryness and reconstituted in 20 μl of 0.1% formic acid for mass spectrometry analysis.

### Histone H1.2 FLAG affinity purification

HEK293T cells were co-transfected in 15 cm format with 3 μg pcDNA4/histone H1.2-3xFLAG with 12 μg of pcDNA4 or with 60 ng pcDNA4/VPR-2xStrep and 11.04 μg pcDNA4 using the PolyJet transfection reagent (SignaGen). Cells were harvested 48 hours after transfection by lysis in cold IP buffer (5 mM Tris-HCl pH 7.4, 150 mM NaCl, 1 mM EDTA, Complete EDTA-free protease inhibitor tablet) with 0.5% NP-40. Debris was pelleted and 40 μL of a 50% slurry of anti-FLAG M2 magnetic beads (Sigma) were added for 2 hours. The sample was eluted by with 100 μg/ml 3xFLAG peptide (Elim Bio) in IP buffer with 0.05% Rapigest SF (Waters).

Affinity purified samples were added to a buffer containing 2M urea, 10 mM ammonium bicarbonate, and 2 mM DTT. Samples were reduced at 60°C for 30 minutes and alkylated with 2 mM iodoacetamide for 30 min at room temperature in the dark.

Trypsin was then added at a 1:100 enzyme:substrate ratio. Samples were digested for 18 hours at 37°C and then desalted using C18 STAGE tips, evaporated to dryness, and then resuspended in 0.1% formic acid for mass spectrometry analysis.

### Denaturing ubiquitin immunoprecipitation

For denaturing ubiquitin immunoprecipitation experiments, HEK293T cells were co-transfected in 6 cm format with 500 ng pcDNA4/histone H1.2-3xFLAG, 500 ng pcDNA/Myc-Ub, 500 ng pMaxGFP, and pcDNA4/TO to bring the total transfected amount to 2 μg. Cells were harvested 48 hours after transfection by pelleting cells, lysing in 150 μL SDS lysis buffer (1% SDS, 50 mM Tris-HCl pH 8.0, 150 mM NaCl) and boiling at 95°C for 10 minutes. Samples were sonicated to sheer DNA and insoluble material was pelleted by centrifugation at 14,000 x *g*. 10 μL of a 50% slurry of anti-FLAG M2 magnetic beads (Sigma) was incubated with the samples for 2 hours at 4°C and then were washed three times with RIPA buffer. Samples were eluted in 30 μL Laemmli sample buffer (Bio-Rad) and boiling at 95°C for 10 minutes. Samples were then separated by gel electrophoresis using a Criterion TGX gel (Bio-RAD), transferred to a nitrocellulose membrane, blocked with either 5% milk or 2% BSA in Tris-buffered sale (TBS) with 0.1% Tween-20, incubated with antibody, and developed with Pierce ECL Western Blotting Substrate (Thermo) and autoradiographic film (Amersham). Antibodies were used at the following concentrations: Myc-HRP (Thermo) at 1:5000, rabbit anti-FLAG (Sigma) at 1:5000, mouse anti-GAPDH (Sigma) at 1:2000, mouse anti-Strep II (Sigma) at 1:2000, goat anti-mouse-HFP (Bio-Rad) at 1:10000, and goat anti-rabbit-HRP (Bio-Rad) at 1:10000.

### Mass spectrometry analysis

Ubiquitination and protein abundance samples were analyzed on a Thermo Scientific LTQ Orbitrap Elite mass spectrometry system equipped with an Easy nLC 1000 uHPLC system interfaced with the mass spectrometry via a Nanoflex II nanoelectrospray source. Samples were injected onto a C18 reverse phase capillary column (75 μm inner diameter x 25 cm, packed with 1.9 μm Reprosil Pur C18-AQ particles). Peptides were then separated by an organic gradient from 5% to 30% in 0.1% formic acid over 112 minutes at a flow rate of 300 nL/min. The mass spectrometry collected data in a data-dependent fashion, collecting one full scan in the Orbitrap at 120,000 resolution followed by 20 collision-induced dissociation MS/MS scans in the dual linear ion trap for the 20 most intense peaks from the full scan. Dynamic exclusion was employed to reject analysis of singly charged species or species for which a charge could not be assigned.

Phosphorylation and AP-MS samples were analyzed on a Thermo Scientific Orbitrap Fusion mass spectrometry system equipped with an Easy nLC 1200 Uhplc system interfaced with the mass spectrometer via a Nanoflex II nanoelectrospray source. The sample capillary columns were used as above. For phosphoproteomics analysis, peptides were separated by an organic gradient from 5% to 30% ACN in 0.1% formic acid over 172 minutes at a flow rate of 300 nL/min. For AP-MS analysis the gradient was over 52 minutes at the same flow rate. In both cases the mass spectrometer collected data in a data-dependent fashion, collecting one full scan in the Orbitrap at 240,000 resolution followed by the maximum number of high energy collision-induced dissociation MS/MS scans that could obtained in the dual linear ion trap within 3 seconds. Dynamic exclusion was enabled for 30 seconds with a repeat count of 1. Charge state screening was applied to reject analysis of singly charged species or species for which the charge could not be assigned.

### Isolation of primary CD4+ T cells

Human T cell isolation and leukoreduction chambers from healthy, anonymous donors were purchased from Blood Centers of the Pacific and processed within 12 hours. Primary CD4+ T cells were harvested by positive selection using a FABian automated enrichment system and CD4 isolation kit (IBA Lifesciences). Isolated T cells were suspended in complete RPMI-1640 medium supplemented with 5 mM HEPES, 2 mM glutamine, 50 μg/ml penicillin/streptomycin, 5 mM nonessential amino acids, 5 mM sodium pyruvate, and 10% fetal bovine serum (Atlanta Biologicals). These cells were immediately stimulated on anti-CD3-coated plates (coated overnight with 10 μg/ml anti-CD3 from Tonbo Biosciences) in the presence of 5 μg/ml soluble anti-CD28 (CD28.2, Tonbo Biosciences). Cells were stimulated for 48 hours prior to electroporation.

### CRIPSR-Cas9 mediated knockout in primary CD4+ T cells

Electroporation was performed using the Amaxa P3 Primary Cell 96-well Nucleofector kit and 4D Nucleofector system (Lonza). Recombinant *S. pyrogenes* Cas9 protein used in this study contains two nuclear localization signal (NLS) peptides that facilitate transport across the nuclear membrane (obtained from the QB3 Macrolab, UC Berkeley). Purified Cas9 protein was stored in 20 mM HEPES pH 7.5, 150 mM KCl, 10% glycerol, and 1 mM TCEP at −80°C. crRNA for each gene were designed by Dharmacon. Each crRNA and the tracrRNA were chemically synthesized (Dharmacon) and suspended in 10 mM Tris-HCl pH 7.4 to generate 160 μM stocks. Cas9 RNPs were prepared fresh for each experiment. crRNA and tracrRNA were first mixed 1:1 and incubated for 30 minutes at 37°C to generate 80 μM crRNA:tracrRNA duplexes. An equal volume of 40 μM *S. pyogenes* Cas9-NLS was slowly added to the crRNA:tracrRNA and incubated for 15 minutes at 37°C to generate 20 μM Cas9 RNPs. For each reaction, approximately 10^5^ T cells were pelleted and resuspended in 20 μL P3 buffer. 4 μL of 20 μM Cas9 RNP mix was added directly to these cells and the entire volume was transferred to a 96-well reaction cuvette. Cells were electroporated using program EH-115 on the Amaxa 4D Nucleofector (Lonza). 80 μL pre-warmed complete RPMI-1640 was added to each well and the cells were allowed to recover for 30 minutes at 37°C. Cells were then restimulated using CD2/CD3/CD28 flow cytometry-compatible stimulation beads (Miltenyi Biotec) in complete RPMI-1640 supplemented with 80 U/ml IL-2-IS (Miltenyi Biotec) and cultured in 96-well V-bottom plates. All subsequent culturing of primary CD4+ T cells was performed in complete RPMI-1640 with 80 U/ml IL-2-IS. Two days post-electroporation media was changed on the cells, and four days post-electroporation cells were split by replica plating into three identical 96-well dishes. Seven days post-electroporation two plates were washed in PBS and banked for genomic DNA purification and Western blot analysis, respectively, and the third plate was used for low MOI HIV-1 infection.

### Low MOI HIV-1 infection of primary CD4+ T cells and flow cytometry analysis

Infected cells were replica plated in technical triplicate in U-bottom 96-well plates, and sufficient HIV-1 NL4-3 Nef-IRES-GFP AS1 reporter virus (Schindler et al., 2003) was added to produce 0.5-2% infection in the first round of infection (according to titration data generated in previous donors) along with sufficient RPMI-164 with IL-2 to bring the total volumne to 200 μL per well. 3, 5, and 7 days post-infected, 100 μL of cells were moved to a separate 96-well plate and fixed in 70 μL of 4% formaldehyde. After collection of each time point, fresh media was added to bring the volume to 200 μL. Flow cytometry was performed on a Becton Dickinson FACSCanto II (BD Biosciences) through the UCSF Laboratory for Cell Analysis. Analysis of flow cytometry data was performed using FlowJo analysis software (version 10.3.0). Quantification was done by first gating the live cell population, followed by gating on the GFP+ cells.

---

**Supplementary Table S1. Global ubiquitination results comparing wild-type HIV-infected vs. mock-infected cells.** (A) MSstats-scored results. (B) Table description.

**Supplementary Table S2. Global phosphorylation results comparing wild-type HIV-infected, vif HIV-infected, and mock-infected cells.** (A) MSstats-scored results. (B) Table description.

**Supplementary Table S3. Global protein abundance results comparing wild-type HIV-infected vs. mock-infected cells.** (A) MSstats-scored results. (B) Table description.

**Supplementary Table S4. Global ubiquitination results comparing wild-type HIV-infected vs. vif/ vpr/ vpu HIV-infected cells.** (A) MSstats-scored results. (B) Table description.

**Supplementary Table S5. Histone H1.2 quantitative AP-MS results with SAINTexpress BFDR < 0.05 comparing cells co-transfected with histone H1.2 and empty vector vs. cells co-transfected with histone H1.2 and VPR.** (A) MSstats-scored results. (B) Table description.

